# A cell cycle-coordinated nuclear compartment for Polymerase II transcription encompasses the earliest gene expression before global genome activation

**DOI:** 10.1101/366468

**Authors:** Yavor Hadzhiev, Haseeb K. Qureshi, Lucy Wheatley, Ledean Cooper, Aleksandra Jasiulewicz, Huy Van Nguyen, Joseph Wragg, Divyasree Poovathumkadavil, Sascha Conic, Sarah Bajan, Attila Sik, György Hutvàgner, Làszlò Tora, Agnieszka Gambus, John S. Fossey, Ferenc Müller

## Abstract

Most metazoan embryos commence development with rapid cleavages without zygotic gene expression and their genome activation is delayed until the mid-blastula transition (MBT). However, a set of genes escape global repression during the extremely fast cell cycles, which lack gap phases and their transcription is activated before the MBT. Here we describe the formation and the spatio-temporal dynamics of a distinct transcription compartment, which encompasses the earliest detectable transcription during the first wave of genome activation. Simultaneous 4D imaging of expression of pri-miR430 and zinc finger genes by a novel, native transcription imaging approach reveals a pair of shared transcription compartments regulated by homolog chromosome organisation. These nuclear compartments carry the majority of nascent RNAs and transcriptionally active Polymerase II, are depleted of compact chromatin and represent the main sites for detectable transcription before MBT. We demonstrate that transcription occurs in the S-phase of the cleavage cycles and that the gradual slowing of these cell cycles are permissive to transcription before global genome activation. We propose that the demonstrated transcription compartment is part of the regulatory architecture of nucleus organisation, and provides a transcriptionally competent, supporting environment to facilitate early escape from the general nuclear repression before global genome activation.

## Introduction

Regulation of transcription underlies the coordination, determination and maintenance of cell identity during organismal development. Nuclear topology and chromatin structure are key factors in the coordination of transcription of genes scattered across the genome (reviewed in^1^). However, the relationship between spatio-temporal dynamics of transcription and the 4D organisation of the nucleus is poorly understood. Genome activation leads to the concerted activation of a large number of genes, which offers a tractable model to address the nuclear topology organisation of dynamic transcription. The earliest stages of development are under the exclusive control of maternally deposited proteins and RNAs, while the embryo remains in a transcriptionally silent state^2^. In externally developing metazoan embryos a series of extremely fast and metasynchronous cell division cycles precede global zygotic genome activation. Genome activation is regulated by a threshold nucleo-cytoplasmic ratio, which is reached at the mid-blastula transition (MBT)^3^ and reflects release from repression by diluted maternal factors such as histones and replication factors^4-7^,. Together with genome activation, simultaneous clearance of maternal RNAs^8, 9^ at MBT overhauls the embryonic transcriptome^10^.

There is accumulating evidence for robust RNA Polymerase II (Pol II) transcription prior to global zygotic genome activation in most animal models^2^. A small group of genes are activated several cell cycles before the MBT and represent the first wave of genome activation^11-13^. The first genes expressed in the zebrafish embryo include microRNAs, which drive the clearance of maternal mRNAs^9^, as well as transcription factors and chromatin binding proteins, which may play a role in the main wave of genome activation^11^. The existence of this first wave of genome activation raises the question of how genes escape the repressive environment before the threshold nucleo-cytoplasmic is reached at MBT. Furthermore, it remains unknown how transcription can occur during the short cell cycles consisting only of S and M-phases. To be able to address these questions, transcription monitoring at single cell resolution is necessary whilst maintaining the developing embryo context. This can only be achieved by *in vivo* imaging. The currently available imaging technologies are based on synthetic transgenic reporters of stem loop RNA-binding proteins fused to fluorescence proteins^14^. This technology allowed monitoring of the dynamics, variation and nuclear topological constraints of gene expression (e.g.^15-18^). However, its limitation is the requirement for transgenic manipulation of each gene of interest. Here we aimed to image native RNAs of endogenous genes without the need to introduce bulky fluorescence proteins by transgenesis. We developed a novel method based on arrays of fluorescently tagged antisense oligonucleotides^19^ for *in vivo* imaging of transcription and transcript accumulation dynamics, which we called <u>MO</u>rpholino <u>VI</u>sualisation of <u>E</u>xpression (MOVIE). Using MOVIE we demonstrate that the earliest gene expression in zebrafish embryos is confined to a unique transcription compartment, which forms during the S-phase of extremely short cleavage stage cell cycles without gap phases. Nuclear organisation of transcription suggests a previously unappreciated nuclear body formation, which is seeded by transcribed loci on pairs of homolog chromosomes and is dependent on cell cycle lengthening. MOVIE enabled investigation of the spatio-temporal dynamics of this transcription compartment, which is a characteristic feature of pre-MBT embryos.MOVIE detects native gene transcription in living zebrafish embryos.

## Results

### MOVIE detects native gene transcription in living zebrafish embryos

We designed a morpholino oligonucleotide (MO) array-based approach to detect 5’ ends of transcripts and their accumulation at endogenous genes in living cells (**Fig. 1a**). We chose the primary transcripts of *miR430* microRNA genes as test for our approach. MiR430 genes are the earliest and highest expressed genes during the first wave of zygotic gene activation^11^ and play a key role in the clearance of maternally deposited mRNA during MBT. We have used cap analysis of gene expression (CAGE) sequencing datasets^20^ to map the transcription start sites of the *miR430* gene cluster. CAGE data revealed at least 8 transcriptional start sites on the reference zebrafish genome (GRCz10), with 6-9 *miR430* genes predicted to be transcribed from each of them (**Suppl. Fig 1a,b**). Two of these promoters contain single nucleotide polymorphisms (SNPs) allowing RNA-seq data to be uniquely mapped, demonstrating that at least 2 *miR430* promoters are used by the embryo (**Suppl. Fig 1c,d**). We hypothesized that multiple promoter use at the locus may facilitate local enrichment of targeted MO signals. We designed a series of three *miR430-* targeting MOs (**Suppl. Table 1**) which upon microinjection into fertilised eggs led to the detection of two distinct transcription foci in nuclei of zebrafish blastula embryos by lightsheet imaging (**Fig. 1a,b**). MO signal followed the developmental stages when *miR430* expression has previously been described^12, 20^, (**Suppl. Fig. 1a**). Expression signals were detected as pairs of foci per nucleus, suggesting detection of *miR430* RNAs from both parental alleles. The signal was RNA dependent, specific to *miR430* RNAs and was lost upon transcription inhibition (**Fig. 1b,c, Suppl. Fig. 1a**). MOs detected *de novo* transcribed nascent RNAs, demonstrated by the loss of fluorescent foci at high stage (10^th^ cell-division), following transcription inhibition at 256-cell stage (8^th^ cell-division) (**Fig. 1c**). MO injection did not affect *miR430* production or development of the embryos (**Suppl. Fig. 2b,c**). We thus demonstrated detection of native and nascent *miR430* primary RNA transcription in normally developing living vertebrate embryos by our novel method, which does not require transgenic manipulation.

**Figure 1.**
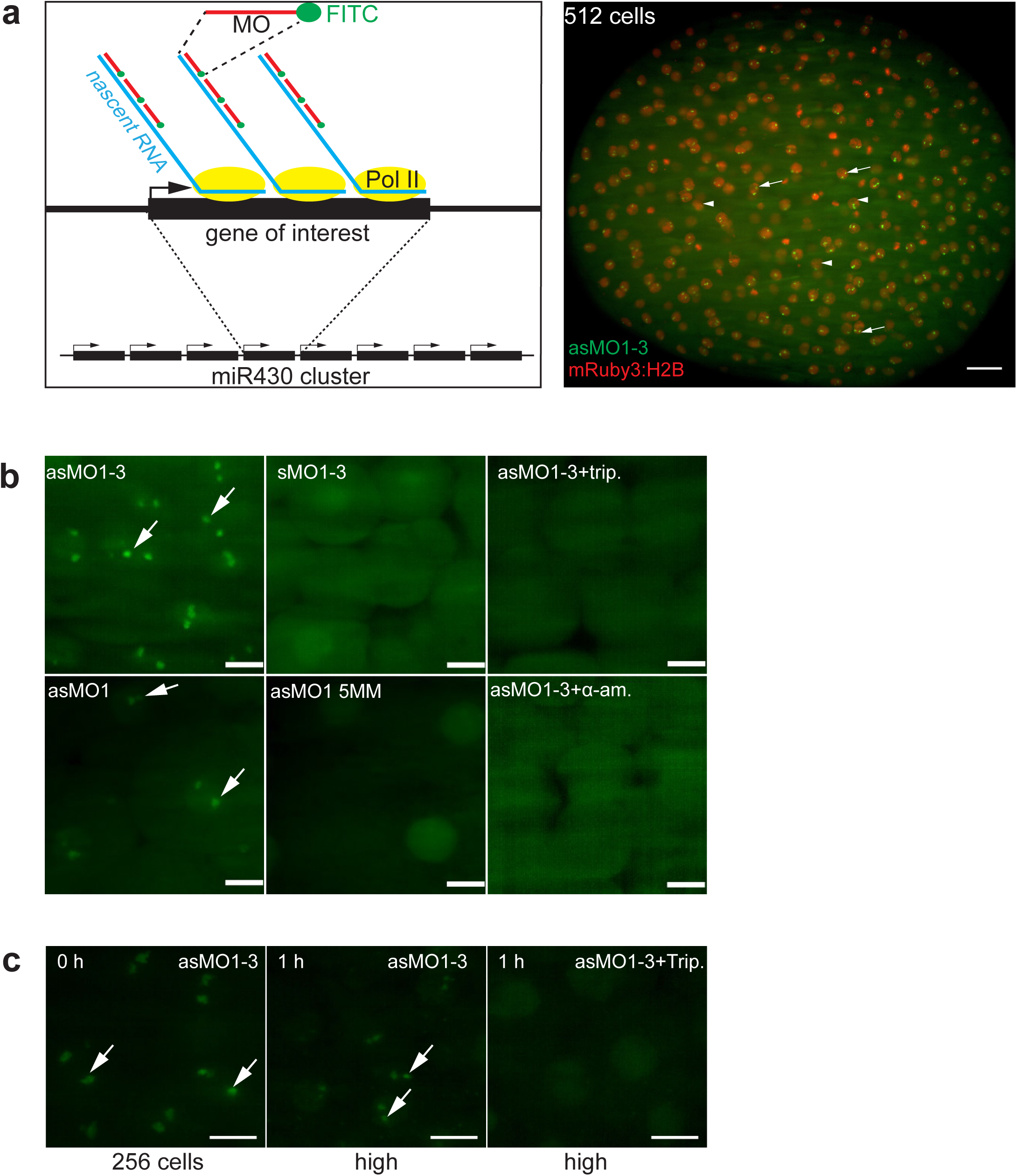
MOVIE detects native transcripts at the site of transcription in living zebrafish embryos. **(a)**) Left: schematic of the detection principle of transcript imaging by MOVIE at the miR430 primary miRNA gene cluster. Right: Image of whole embryo, imaged in vivo by lightsheet microscopy, injected with three antisense, FITC labelled morpholinos (MO) targeting miR430 priRNA (green) and mRuby3:H2B fusion protein as nuclear marker (red). Arrows point at signal accumulation, appearing as double green punctae within the nuclei, labelled by mRuby3:H2B (arrowheads). **(b)**) Representative images of live embryos, imaged in vivo at high/oblong by lightsheet microscopy, injected at zygote stage with triple and single sense, antisense and mismatch containing, FITC labelled morpholinos and transcription blockers (α-amanitin or triptolide, n=6 embryos each). **(c)** Transcript signal detection before and after 1-hour triptolide treatment started at 256-cell stage (0 h), (n=6 embryos each). Stages of embryos as indicated in panels. Abbreviations: trip, triptolide, as, antisense, s, sense, MO, morpholino oligonucleotide (where number depicts specific MO, for details see **Suppl. Table 1**), h, hours after treatment. Scale bars: **a** 50 μm **b-c** 10 μm.

### Transcription of *miR430* is metasynchronous and follows the cleavage stage cell cycle periodicity

Zebrafish cleavage stages are characterised by metasynchronous cell cycles in which synthesis and mitosis phases are cycling without gap phase^3, 21-23^. To understand the temporal relationship between *miR430* transcription and the fast cell cycles of cleavage stages we monitored *miR430* expression by 4D time lapse imaging with MOVIE and simultaneously monitored cell cycles using the nuclear marker mRuby3:H2B (**Fig. 2a, Suppl. Movie 1**). Quantification of the *miR430* transcript detection led to three main observations. Firstly, *miR430* expression was first detected at 64-cell stage matching previous transcriptome analyses^11, 20, 24^, (**Suppl. Fig 1a**) and was present until late epiboly stages in diminishing numbers of cells. The number of cells with *miR430* expression gradually increased during cleavage stages, potentially explaining the increase in genome-wide *miR430* expression quantification data from whole embryos (**Fig. 2b**, **Suppl Fig. 2a**). By the 512 cell stage all detectable nuclei show miR430 expression however, due to the loss of synchrony of this cell cycle the total number of cells expressing at any given time peaks at around 60% (**Fig. 2c**). Thirdly, the temporal distribution of the start and extent of the MO signal showed gradually expanding asynchrony (**Fig. 2c**), which is in line with the previously described^25^ sequentially increasing length of pre-MBT cell cycles after cycle 6 (64-cell stage). Transcriptional activity in these cell cycles also followed a spatial wave-like pattern running across the animal pole (**Suppl. Movie 2**). There was no obvious allelic (parental) bias observed with most cells showing two foci of activity (**Fig. 5g**). Thus transcription of *miR430* genes first detected at the 64-cell stage, does not appear to respond to a precise developmental stage timer co-ordinately. Instead, miR430 expression appears in sequentially increasing number of nuclei and appears to be coordinated with the overall timing of metasynchronous cell cycles.

**Figure 2.**
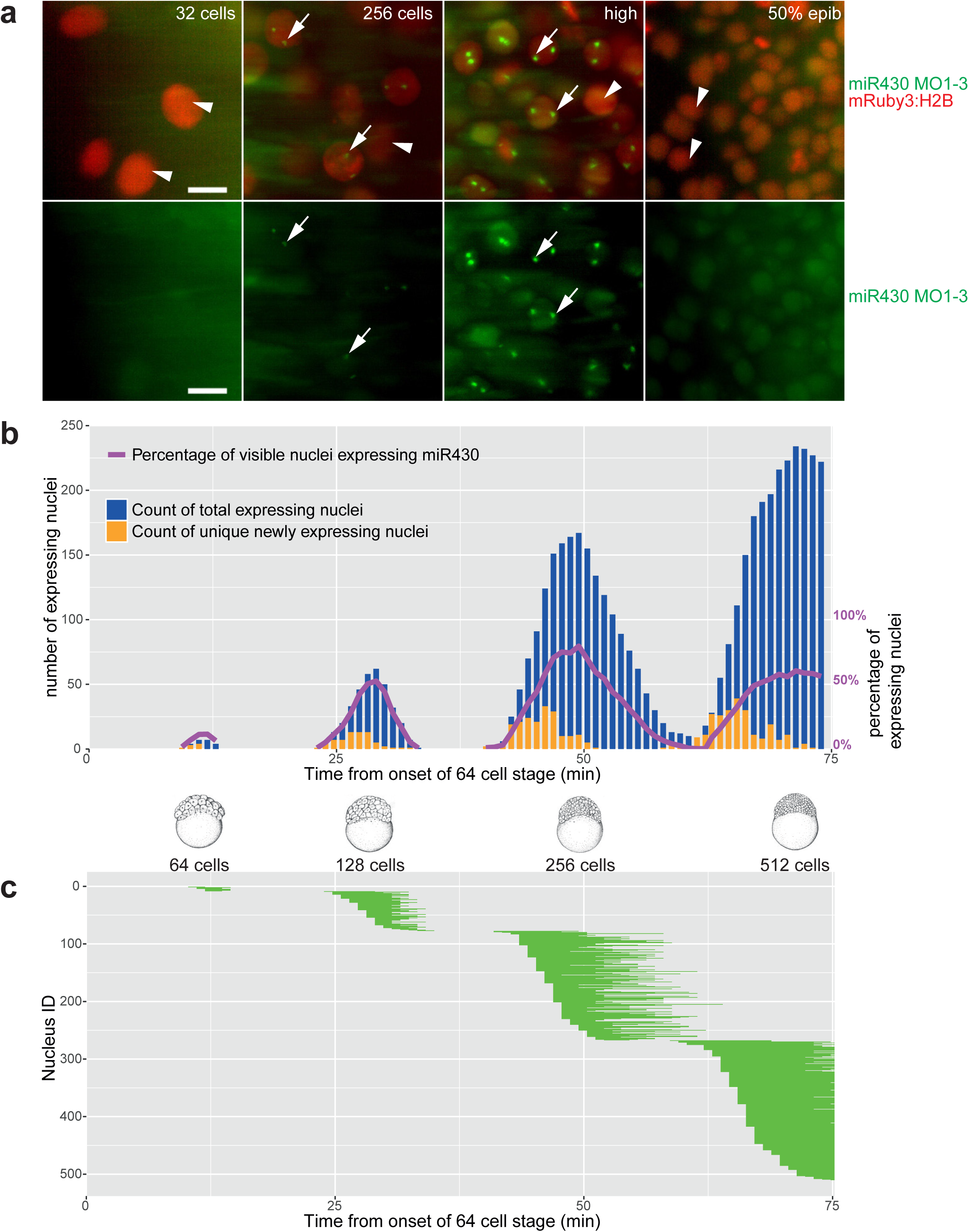
Mapping of gene expression dynamics at cellular resolution indicates that metasynchrony and periodicity follows the short cell cycle phases. **(a)**) Representative examples of simultaneous detection of miR430 expression (arrows, green signal of FITC-labelled MO) with nuclear division cycles (arrowhead, mRuby3:H2B in red) during early development at stages as indicated. **(b)**) Timing of the commencement of gene expression (orange) and number of nuclei (green) showing expression activity, total and unique miR430 expressing nuclei per frame. Stages of development are indicated at the bottom of the panels, purple line represent percentage of nuclei expressing miR430 at the time and stages as indicated. **(c)** Expression span of individual nuclei over time. Each horizontal bar represents the miR=430 expression time of a single nucleus. Chart x-axis matches that in **b**. Each bar represents cumulative signal from up to 2 foci per nucleus. The frame rate (51s) was used as reference for the elapsed time in **b-c** Abbreviations: 50% epib, 50% epiboly. Scale bars: **a** 10 μm

### Transcription and transcript accumulation of *miR430* occurs in the S-phase of permissive, elongating cell cycles

The single-cell resolution of MOVIE also allowed us to address the temporal dynamics of each cell’s transcriptional activity during metasynchronous cell cycles. We used Tg(Xla.Eef1a1:h2b-mRFP1) transgenic embryos^26^ or mRuby3:H2B microinjected protein to analyse cell cycle length. Neither the red fluorescence-tagged H2B, nor MO injection affected the cell cycle (**Suppl. Fig. 3b,c**) allowing the accurate monitoring the relationship between cell cycle and transcription. Cell cycle length was measured by calculating the time between two anaphases. Elongation of these cell cycles, which was previously demonstrated to be due the elongation of the S-phase^21^ was accompanied by elongation of miR430 signal detection (**Fig. 3a,b**). Cell cycle length varies among cells of a single cleavage stage cycle^21^ and we asked whether a correlation between *miR430* expression length and cell cycle length can also be detected among individual cells within a single cleavage cycle. As shown in **Fig. 3c**, positive correlation was observed between both cell cycle length and S-phase length but not mitosis length versus *miR430* expression. Next, we asked when *miR430* expression occurs within the alternating M and S-phases of the pre-MBT cell cycles. By using morphology distribution and entropy of mRuby3:H2B pixels as indicators for chromatin compaction we segmented the cell cycle into its mitotic phases (**Suppl. Fig. 3d**) measured their length (**Fig. 3a,d**). We detected the start of *miR430* activity universally during the S-phase of the pre-MBT cell cycles but with varying delay (**Fig. 3d**), while lowest levels of activity are potentially missed by the imaging approach.

**Figure 3.**
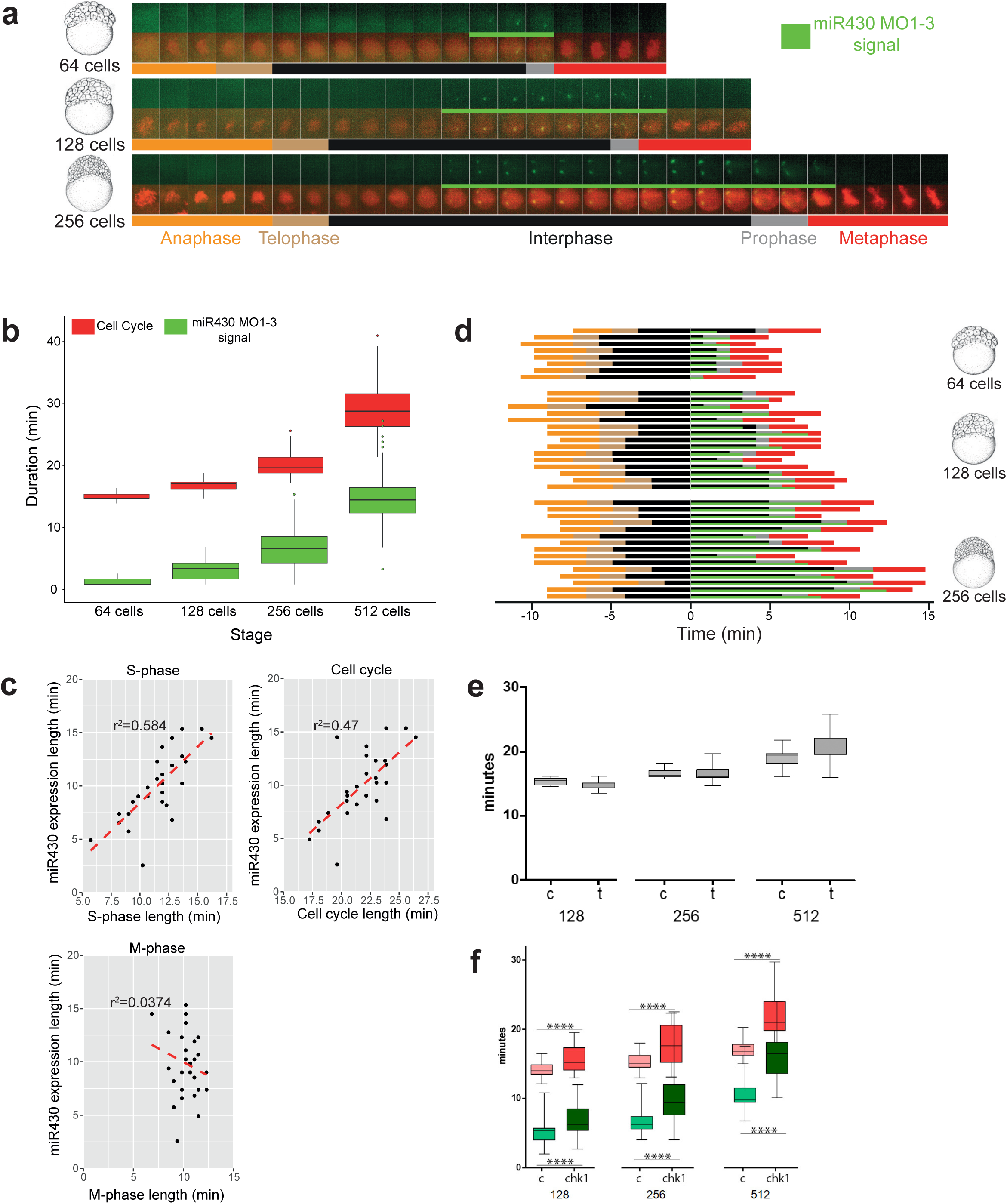
The S-phases of cleavage stage cell cycles are permissive to transcription and transcript accumulation. **(a)** Cell cycle length (anaphase to anaphase detected by mRuby3:H2B in red) and miR430 expression (MO signal in green) during early blastula stages. Frames were acquired in 49s intervals. **(b)** Box plot quantification of elongation of miR430 signal (green) alongside cell cycle length (red). Data from 2 embryos are calculated. Total number of nuclei (cell cycle): 64-cell=17, 128-cell=60, 256-cell=62. All groups were significantly different from one another (p<0.001) (One way ANOVA with multiple comparisons). **(c)** Correlations of cell cycle, S-phase and M-phase length with miR430 expression length, at 256-cell stage (2 embryos, 15 nuclei each). **(d)** Cell cycle phase segmentation as shown in **a**. Horizontal bars represent a nucleus each, they are aligned to onset of detection of miR430 expression and ordered according to length of miR430 expression and cell cycle length. X-axis shows time in minutes. **(e)** Effect of triptolide treatment on cleavage stage cell cycle length. Lightsheet microscopic images for nuclear pattern of mRuby3:H2B were analysed to assess cell cycle length and data merged from 4 embryos, 15 nuclei each. Water-injected controls (c) and triptolide-treated (t) are shown at 128, 256 and 512-cell stage respectively. No statistical difference was found between c and t by Mann Whitney U test. **(f)** Relationship between the length of cell cycle measured by mRuby3:H2B pattern (red) and miR430 activity measured by MO signal (green) in control RNA and chk1 mRNA injected embryos. Data was analysed from 4 embryos, 15 nuclei each at stages as indicated below the charts. Statistical significance was measured by Mann Whitney U test, p< 0.001, whiskers show max and min values.

An important observation from the cell cycle phase segmentation analysis was the detection of MO signals, albeit in reduced volume, into the prophase and even extending into metaphase (**Fig. 3a,d**) either suggesting transcription during M-phase or transcript retention at the transcription site.

Spatio-temporal coupling of replication and transcription at the main wave of zygotic genome activation has previously been established^23, 27-30^, yet this has not been explored during the little studied first wave of genome activation. Our observation that the temporal extension of *miR430* activity matches that of the corresponding cell cycle lengthening suggests that transcriptional activity is co-regulated with cell cycle elongation during cleavage stages. To test if transcription triggers elongation of the length of cell cycles at these early stages we have blocked transcription initiation by triptolide. Embryos with blocked transcription failed to epibolise as shown previously^31^ (**Suppl. Fig. 4a**). No significant change was observed in cell cycle length, suggesting that cell cycle elongation in cleavage stage embryos is not explained by transcriptional activation (**Fig. 3e**). In contrast, when we have forced precocious checkpoint kinase 1 (Chk1) activation by injecting synthetic *chk1* mRNA^32, 33^, we detected significant elongation of the cell cycle in cleavage stage embryos and corresponding elongation of miR430 MO signal (**Fig. 3f**, **Suppl. Fig. 4b**) in every cleavage cycle measured. Thus, cell cycle elongation is permissive for *miR430* transcription and transcript accumulation in the sequentially expanding S-phases during cleavage stages in zebrafish.

### First wave of gene expression is confined to a primary transcription compartment shared by genes

The observation of a pair of *pre-miR430* transcription accumulation foci raises the question of the nature and dynamics of this compartment in relation to other transcriptional activities during the first wave of genome activation. We used an antibody that recognises phosphorylated serine-2 in the C-terminal domain (CTD) of the large subunit of RNA Polymerase II (Pol II Ser2P) as a marker of transcription elongation^34^. This revealed two large foci, similar to *miR430* in pre-MBT stages that expand to broadly distributed irregular speckles after the main wave of global genome activation, likely reflecting the large expansion of transcription at thousands of genes^10, 12^(**Fig. 4a**). We used an *in vivo* approach to simultaneously detect *mir430* and Pol II SerP2 activities. A single *miR430* targeting MO, which was previously observed to be sufficient to detect *miR430* activity alone (**Fig. 1b**), was chosen for labelling with Cy5 (Suppl. material) and co-injected with labelled Fab fragments against Pol II Ser2P^35^ into embryos to image by confocal microscopy *in vivo*. Colocalisation pattern was observed between Pol II Ser2P to miR430 MOs in live embryos, indicating that these newly described miR430-containing domains are transcriptionally highly active (**Fig. 4b**) at pre-MBT stages and appear to be the main site of Pol II activity at these stages. The colocalisation of Pol II SerP2 with *miR430* transcripts was also verified by immunostaining (**Fig. 4b**). To further characterise cleavage stage transcription sites by an independent approach, we used a nucleotide analogue incorporation assay^36^ and visualised nascent transcripts in pre-MBT embryos with and without Pol II Ser2P localisation. This showed colocalisation of the majority of nascent RNAs together with Pol II Ser2P (**Fig. 4c**). Furthermore, immunohistochemical detection of H3K79me2, an indicator of transcriptionally active chromatin^37^ at the site of Pol II Ser2P provides additional evidence for the majority of transcriptional activity being confined to two transcription compartments at these stages of development (**Fig. 4d**). A 200kbp BAC probe containing the *miR430* locus was detectable in nuclei as large but distinct nuclear territories and colocalised with the main site of nascent RNA detection (**Fig. 4e**) and Pol II Ser2P (data not shown). This supports the suggestion that *miR430* activity marks the transcriptional compartment, which appears as the main site of early transcription and a characteristic of the first wave of genome activation.

**Figure 4.**
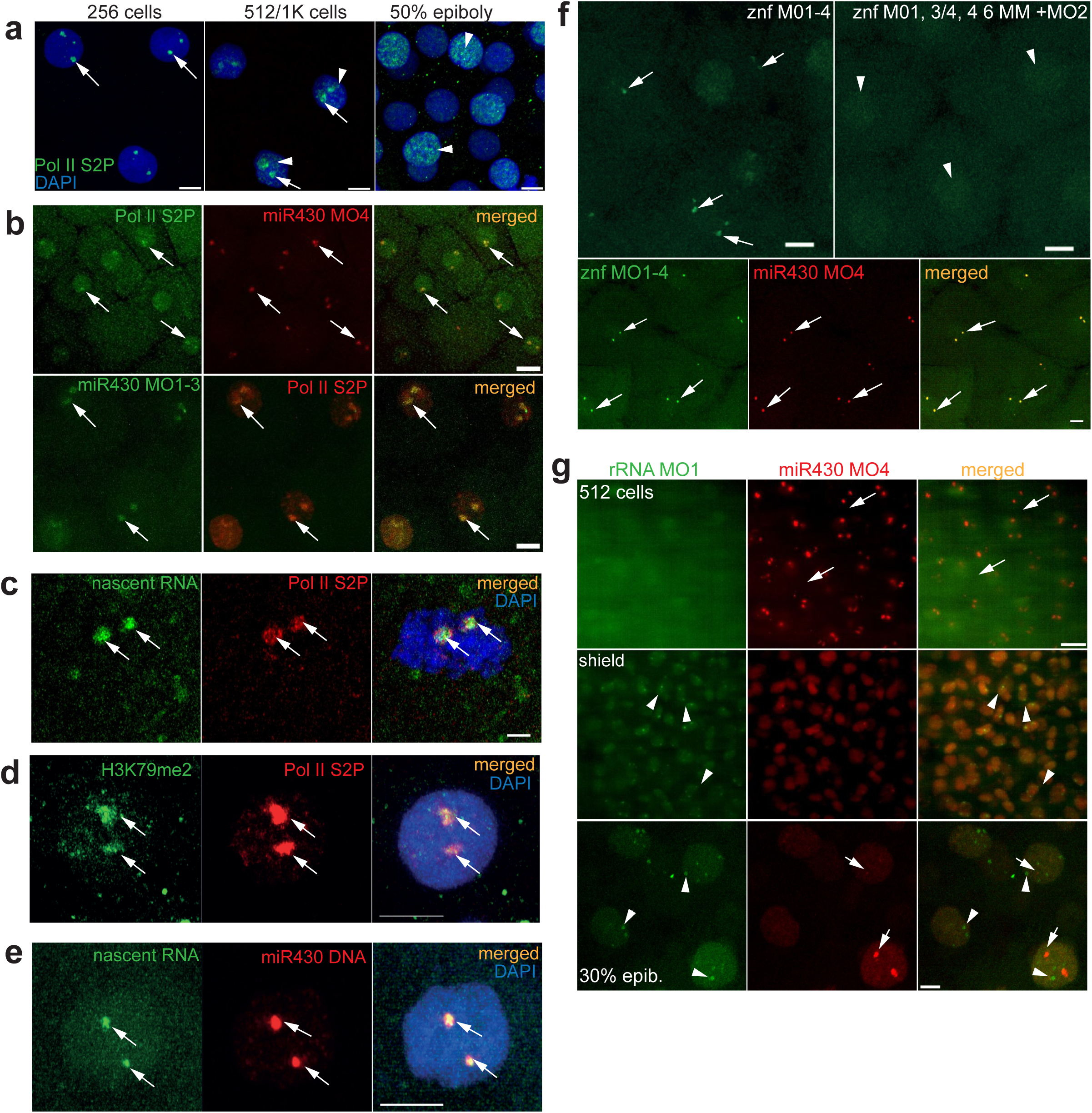
First wave transcription is confined into a nuclear compartment. **(a)** Immunohistochemistry of Pol II S2P (green) at stages as indicated of Pol II S2P at two main foci at pre-MBT (arrows) and broader distribution (arrowheads) at relevant stages. Nuclei are stained with DARI (blue). **(b)** Top panel: live imaging of embryos at 256-cell stage, injected with anti Pol II Ser2R fab fragments, labelled with Alexa Fluor 488 (green) and Cy5-labelled miR430 targeting MO4 (red). Green and red signals colocalise in all nuclei, where both signals are detected (n=3 embryos). Bottom panel: immunohistochemistry of Pol II S2P (red) and miR430 targeting MO1-3 (green) at high stage; green and red signals colocalise in all nuclei (3 embryos). **(c)** Immunohistochemistry of Pol II S2P (red) and nascent RNA detection (green) by ethynyl-uridine incorporation and subsequent Alexa Fluor 488 tagging of embryos at 256-cell stage. Green and red signals colocalise in all nuclei (n=5). **(d)** Immunohistochemistry for H3K79me2 (green) and Pol II Ser2R (red) and merged image with DARI in blue, performed on 512-cell stage embryos. The third panel shows these signals merged with DARI in blue (8 nuclei in 3 embryos). **(e)** Staining for the miR430 locus (red) in combination with nascent RNA detection (green) by ethynyl-uridine incorporation in embryos at 512-cell stage. Merged panel shows DARI in blue (n=16 nuclei). **(f)** Top panels: live detection of activity of a cluster of znf genes at 512-cell stage by fluorescein-labelled MOs, injected with mix of targeting (left) or mismatch-containing control MOs (right) (n=3 embryos each). Bottom panels: embryo co-injected with miR430 MO4 (red) and znf targeting MOs (green) at 256-cell stage. Green and red signals colocalised in all nuclei (n=6 embryos). In panels b-f arrows point at transcription foci. Arrows point at transcription foci (**b-f**) **(g)** Top: Lightsheet images of embryo (at 512-cell and shield stages) co-injected with MOs targeting the 5’ETS of the 45S ribosomal RNA precursor (green) and miR430 MO4 (red). Bottom: high magnification confocal Airyscan image of 30% epiboly embryo injected with the same reagents. miR430 MO4 signal accumulation (arrows) and rRNA_5'ETS_MO1 (arrowheads) are indicated. Abbreviations: znf, zinc finger gene; Pol II S2P, RNA polymerase II serine 2 phosphorylated; rRNA, ribosomal RNA; ETS, external transcribed spacer, Scale bars: **a**, 10 μm; **b** top 20 μm, bottom 10 μm; **c**, 2 μm **d-e**, 5 μm; **f**, 10 μm; **g**, top 20 μm bottom 5 μm.

Next, we asked whether expression of other genes besides *miR430*, contribute to the formation of this transcription compartment. We queried a partially annotated set of *zinc finger (znf)* genes, which consist of over 350 copies localised on the same chromosome as *mir430* (chr4)^12^ and using CAGE-seq data we found evidence for their expression during the first wave of genome activation (**Suppl. Fig. 5a**). From these, 25 *znf* genes are highly conserved (**Suppl. Fig 5b**), highly active at pre-MBT and early post-MBT and scattered into 37 Mbp region, immediately downstream of the *miR430* gene cluster on chromosome 4. The pre-MBT activity of *znf* genes suggested them as candidates for transcription visualisation and we designed MO sequences to the highly conserved 5’ end of their transcripts (**Suppl. Fig 5b**, **Suppl. Table 1**). A series of quadruple MOs, which together targeted all 25 *znfs* (**Suppl. Fig 5c**) were injected into embryos and led to specific detection of *znf* gene activity in two foci similar to *miR430* during cleavage and MBT stages (**Fig. 4f**). Using two colour labelling of MOs for the co-detection of fluorescein-tagged *znf* MOs and Cy5-tagged miR430 MO4, we demonstrated colocalisation of these two sets of RNAs in a pair of shared transcription compartments (**Fig. 4f, Suppl. Movie 3**).

Next we asked about the relationship of the observed transcription compartment with other compartment forming nuclear transcriptional activities. Four out of five loci of 45S rDNA, which encode Pol I-transcribed ribosomal RNAs (rRNA) reside on chromosome 4^38^. We asked whether nucleoli forming rRNAs can also be visualised similarly to Pol II transcribed genes on the same chromosome. Using MOVIE we found that rRNA transcripts can be detected, however only after the main wave of genome activation at MBT, with low signals at 30% epiboly and frequent activity at shield stages (**Fig. 4g**) which is in line with previously described rRNA expression patterns^36, 38^). At post-MBT stages *miR430* transcripts were still detected in a mosaic fashion (**Fig. 4g**) and allowed subnuclear localisation analysis of both RNAs when both gene products were detected in the same cells. We thus used dual MO labelling and compared *miR430* RNA and rRNA distribution. At 30% epiboly stage their accumulation was clearly distinctly localised (**Fig. 4g** bottom row). This result demonstrates that the transcription compartment containing Pol II gene products is independent from the nucleolus-forming RNA compartment despite their loci residing on the same chromosome.

### Relationship of the transcription compartment with the mitotic nuclear architecture

Our results, obtained by several imaging tools, demonstrate the formation and prominence of a major transcription compartment, characterised by localised Pol II Ser2P enrichment and high concentration of newly synthesized nascent RNAs of several genes from at least two sets of gene families. Thus, it may fulfil several of the criteria for what previously has been described as a phase separated nuclear body (reviewed in^39, 40^). If the transcription compartment is a phase separated nuclear structure, local depletion of chromatin is expected in the compartment as has recently seen during formation of transcriptional microenvironments during the main wave of genome activation^41^. To test this, we have analysed nuclear structure in pre-MBT embryos by simultaneous MOVIE imaging of *miR430* RNA together with labelled H2B and labelled Proliferating Nuclear Antigen (PCNA), which is broadly localised to DNA in the nucleus during S-phase of cleavage stages in zebrafish^42^. Reduction of both chromatin-associated proteins was observed at the site of *miR430* RNA accumulation and this depletion correlated with the dynamics of RNA detection (**Fig. 5a**). This lack of chromatin in a compartment was also seen without MO injection, or in embryos where endogenous Pol II Ser2P was detected without MO injection (**Fig. 5b** and data not shown). Thus, we describe a nuclear transcription compartment, with coding and non-coding Pol II transcribed nascent RNAs, large quantities of transcriptionally active Pol II and local depletion of compact chromatin, which together suggest the formation of a phase separated transcription body in early embryos.

**Figure 5.**
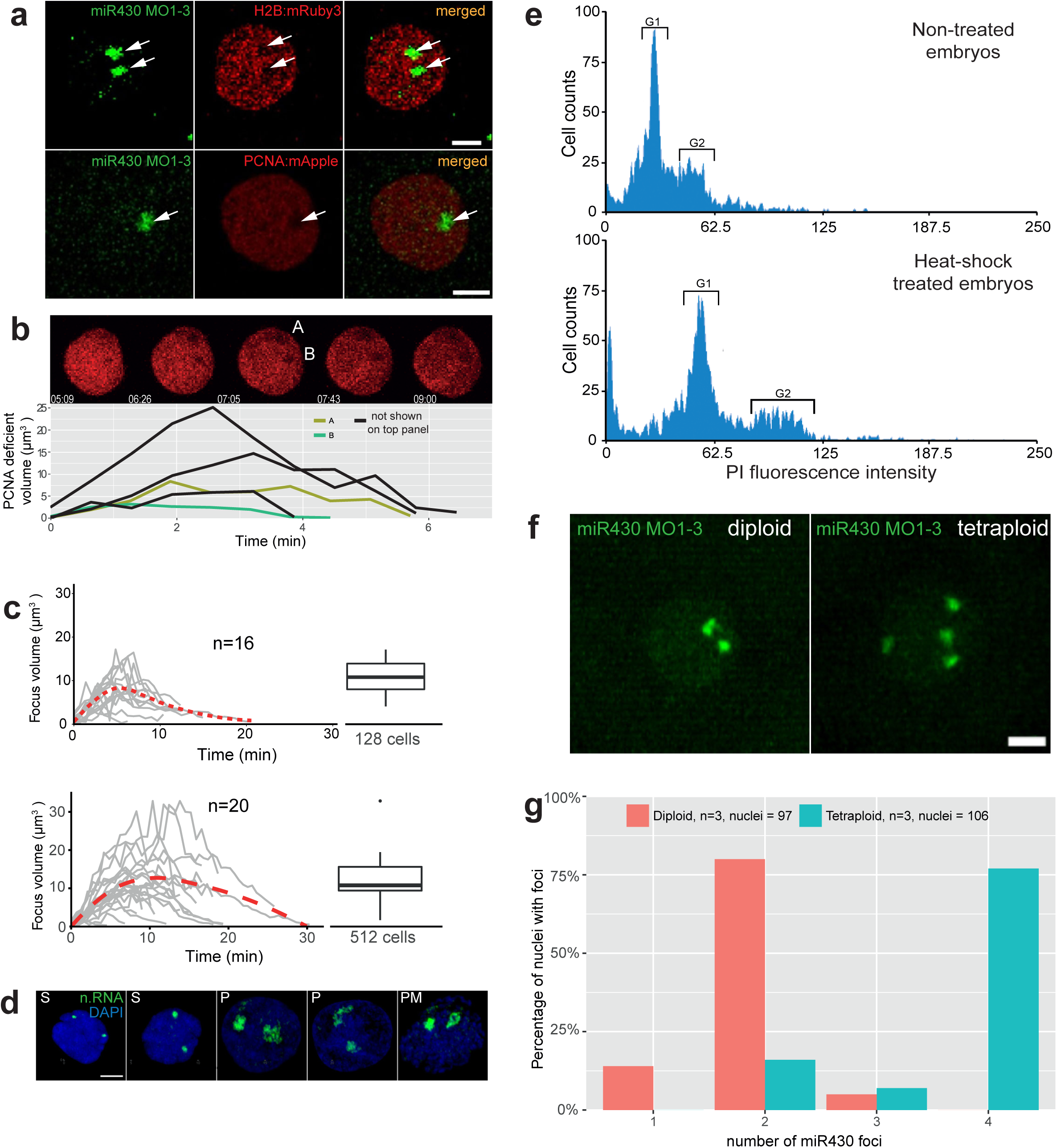
Relationship of the transcription compartment with the mitotic nuclear architecture. (**a**) Confocal Airyscan image of 512-cell stage embryos co-injected with miR430 MOs (green) and either mRuby3:H2B (top) or mApple:PCNA (bottom) fusion proteins (red). Arrows indicate low density chromatin at the site of miR430 activity. **(b)** Timelapse of 512-cell stage nucleus, injected with mApple:PCNA fusion proteins (red). Image shows individual z-slices from within the nucleus exhibiting two PCNA deficient regions. Graph below shows the evolution of the individual depicted PCNA deficient regions over time (A & B), in addition to deficient regions from other 512-cell stage nuclei from the same embryo, detected using spot detection plugin in Icy Bioimage. Time indicates duration from first detection of PCNA signal within the cell cycle in seconds and minutes. **(c)** Charts representing the transcription focus volume over time based on miR430 MO signal, measured using spot detection plugin Icy Bioimage software, top panel: 128-cell stage, bottom: 512-cell stage. Boxplots on the right show variation in peak volume. **(d)** Nascent RNA detection (green) by ethynyl-uridine incorporation and subsequent Alexa fluor 488 tagging of embryos fixed in 4 min intervals within 512-cells stage, demonstrating different stages of the transcription compartment structure during the cell cycle **(e)** DNA content of 12-somite stage embryos untreated and heat-shocked was determined by propidium iodide DNA staining analysis. FACS traces show a shift in the position of the Gap phase 1 (G1) and Gap phase 2 (G2) peaks (marked), between the heat-shock untreated and treated groups, in accordance with a doubling in DNA content in the heat-shocked group. **(f)** Representative images of miR430 MO-injected diploid and induced polypoid 256-cell stage embryos (green) **(g)** Bar chart showing comparison of miR430 foci number between diploid and tetraploid embryos. Abbreviations: n.RNA, nascent RNA; S, S-phase; P prophase; PM prometaphase; Scale bar, 5 μm.

Next we addressed the temporal dynamics of the nuclear compartment in relation to cleavage stage cell cycles. Measurement of miR430 MO signal and chromatin depletion volumes indicated highly dynamic formation, maintenance and disintegration of this compartment within a cell cycle (**Fig. 5b,c Suppl. Fig. 6a, Suppl. Movie 4**). The transcription compartment has a characteristic volume dynamic, which shows gradual increase at both 128 and 512-cell stages peaking approximately at 16 μm^3^. At 512-cell stage the maximum volume shows an extended plateau in comparison to 128-cell stage, which correlates with the extended cell cycle (**Fig. 5c, Suppl. Movie 5**). The transcription compartment and its dynamics were also confirmed independently without MO injection in fixed embryos using nascent RNA detection by EU incorporation (**Fig. 5d**).

The paired nature of the transcription compartments suggests that their formation is seeded by anti-paired homolog chromosomal loci. To test the role of chromosomal organisation in forming transcription compartments, we generated tetraploid embryos by blocking centrosome separation during the first cell division using heat shock^43, 44^ and verified their ploidy by FACS (**Fig. 5e,f**). We counted *miR430* activity foci and found that the overall majority matched diploid allele count, whereas the number of foci was double that of diploid’s in tetraploid embryos (**Fig. 5g, Suppl. Movie 6**). This finding suggests that the transcription compartments are regulated by mitotic organisation of homolog chromosomes during cleavage stages. Interestingly, tetraploids, which have double the nucleo-cytoplasmic ratio and initiate MBT one cell cycle earlier to normal diploid wild types^3^ did not activate *miR430* expression earlier than diploids, which argues against a nucleo-cytoplasmic ratio threshold being the trigger for their gene expression before MBT (**Suppl. Fig. 4c**).

## Discussion

Here we describe a unique transcription compartment, the formation of which is characteristic to the first wave of gene expression in the embryo. Our results suggest that pre-MBT transcription escapes the global repressive effect of diluted maternal factors via the formation of a dynamic transcription compartment, where most detectable transcription from at least two gene families takes place. The observation that the majority of nascent RNAs and Pol II Ser2P colocalise with local depletion of chromatin indicates the formation of a primordial nuclear body as the major site of detectable transcriptional activity prior to global genome activation at MBT.

The temporal dynamics of this transcription compartment follows and is co-regulated with the metasynchronous cell divisions both of which gradually elongate and increasingly lose synchrony at each cell cycle. This argues against long held views on MBT as a watershed stage; instead supporting a gradual process model for genome activation, previously proposed, based on gradually elongating cell cycles^45^. By manipulating transcription and cell cycle progression we demonstrated that these sequentially expanding cell cycles are permissive for transcription and transcript accumulation and that the majority of transcriptional activity is during S-phase. Yet, it is unclear whether the observed presence of transcripts during prophase and prometaphase are remnants of nascent RNAs or indicate *de novo* transcription. It is notable, that Pol II Ser2P is still present at prophase, potentially reflecting transcriptional activity in M-phase however, it is unclear whether this reflects elongating polymerase as residues of Pol II Ser2P can still be detected in α-amanitin and triptolide injected embryos (data not shown).

The mechanisms that allow pre-MBT transcription are likely different from that underlying the main wave of genome activation. Several mechanisms were suggested to underlie transcription activation in early embryos. These include a nucleo-cytoplasmic (NC) ratio threshold acting via titration of maternal factors^4^, a developmental timer acting via delayed access to transcriptional activators regulated by time-limited translation of maternal mRNAs (e.g.^10^) or availability of threshold amount of DNA template (reviewed in^8, 46^). In our experiments NC ratio does not appear to regulate the earliest of gene activation as manipulation of nuclear content by inducing polyploidy did not have a marked effect on the timing of when expression is first detected. Furthermore, our observations on pre-MBT transcription detection at the single cell level and with specific loci also exclude the model, which argues for lack of detectable transcription due to limiting amount of DNA template. Instead, our data suggests alternative mechanisms, which could include delayed availability of transcriptional activators, for example by the control of maternal mRNA translation.

By detection of specific chromosomal loci and their RNA products, as well as by ploidy manipulation, we demonstrated that these nuclear bodies are seeded by DNA and that the candidate loci for coordinating these nuclear compartments reside on anti-paired homologs of chromosome 4. While our rRNAs imaging data suggests separation of Pol II and nucleolus forming rRNAs compartments, future studies will be necessary to investigate the relationship between the transcription compartment described here and other nuclear bodies, such as Cajal body and histone locus body during early zebrafish development^36^.

The formation of the transcription compartments may serve several purposes in the uniquely fast cell division cycles, which precede global genome activation. Firstly, it may allow for efficient transcription by concentrating the rate limiting transcription apparatus into a confined nuclear space, at a time, when activators and transcription machinery components are likely rate limiting by delayed translation of maternal stockpiles of their mRNAs^24, 47^. Secondly, restricted localisation of transcription into a compartment may indicate an as yet unexplored mechanism for facilitating the escape from the general repressive environment of the nucleus, which is only removed from the rest of the genome by reaching a certain NC ratio. However, initial observations suggest that the first wave of transcription is not affected by similar titration of maternal repressive factors as the rest of the genome, as indicated by unaffected initiation of transcription in tetraploids, where the NC ratio is doubled. Thirdly, it may provide spatial and temporal separation of replication and transcription- the two potentially conflicting genome activities occurring in the short S-phases without gap^48^.

The compartmentalisation of the first wave of genome activation is likely a general phenomenon and not unique to zebrafish. *In situ* detection of large Pol II Ser2P and Pol II pSer5 punctae in *Drosophila* embryos during and before global genome activation, has been documented^49-52^ and may explain the first wave of genome activation not only in *Drosophila* but more generally among externally developing animal models. Due to the simplicity of the MOVIE approach, testing this hypothesis in early embryos of other animal models will be feasible. Utilising multicopy gene product detection, MOVIE will be particularly useful to study RNA accumulation in nuclear bodies. Taken together, our *in vivo* monitoring of nuclear architecture organisation into transcription compartments contributes to the growing body of evidence supporting the notion of suprachromosomal organisation in the nucleus, serving not only structural, but also gene regulatory function e.g.^53-55^ and highlights the early embryo as a tractable model system to study the interface between transcription and nuclear organisation.

## Acknowledgements

This work was supported by a Wellcome Trust Investigator award and BBSRC project grant to FM, NEURAM and VISGEN projects of H2020 by the European Commission to FM, AS, HQ & JF. BBSRC MIBTP doctoral training programme to LC, and European Research Council Advanced grant (ERC-2013-340551, Birtoaction) to LT. We thank Gene-tools for offering Morpholino reagents, the COMPARE imaging facility at the University of Birmingham, Dr Aditi Kanhere and Anja Baranasic for advice in computational analyses, Sarah Baxendale for providing transgenic lines and Phil Zegerman for *chk1* and replication factor expression vectors, Marissa Williams and Glen Reid for assistance with the ddPCR and Piotr Balwierz for comments on the manuscript.

## Methods and materials

### Characterisation of *miR430* locus

Identification of predicted multiple promoter sites at the *miR430* locus were determined from CAGE-seq data^20^ mapped to zebrafish assembly GRCz10 by Tophat^56^ allowing up to 80 multi-mapped reads. Start sites of transcription were determined by usage of CAGEr software^57^. Annotation of promoter regions (Supplementary Figure 1) were defined as 400 bases upstream and 100 bases downstream of the detected TSS with *pre-miR430* gene annotations determined from Ensembl database with addition of 2 pre-*miR430* c-subtype sequences found by BLAT search, completing the a-c-b triplet structure^58^.

Detection of unique SNPs in *miR430* promoter sequences was achieved by multiple sequence alignment with Clustal Omega^59^. Thus allowing detection of nascent RNA^11^ reads aligning to two individual promoters in the *miR430* cluster by unique mapping with STAR aligner^60^. Expression of *miR430* during development is demonstrated both by CAGE–seq^20^ and RNA-seq^12^ spanning 19 developmental time points, capturing expression both pre and post-MBT (Supplementary Figure 2d).

### Morpholino oligonucleotides design for detecting transcription of zinc finger genes on chromosome 4

Targeting morpholino oligonucleotides were designed specifically to visualise transcription of the early zygotic, most highly, expressed zinc finger genes on chromosome 4 as determined from CAGE-seq dataset^20^. Twenty three such genes with a fully annotated transcript were identified. The 5’ UTRs of these genes with 50 bp up- and downstream flanking sequence were used for conservation assessment via multiple sequence alignment using MUSCLE alignment algorithm^61^ and subsequent visualizations of inherent similarity were achieved with JalView^62^. Consensus sequence was generated and used for morpholino design, targeting conserved regions in the 5’UTRs. Altogether 4 morpholino were designed (Supplementary Figure 5b).

To identify all potential *znf* genes on chromosome 4 targeted by the designed morpholino oligonucleotides, a BLAST search against Ensembl version 92 cDNA database was performed. A transcript was considered as potential target by a morpholino if at least a 20 bp match with 84% identity was found. Intersections were calculated from the identified potential target genes list for each morpholino by VennMaster^63^ and quad venn diagram graphic (Supplementary Figure 5c) was generated with VennDiagram CRAN R package^64^.

### Microinjection of morpholinos fusion proteins and antibodies

Single-cell stage embryos were injected with 2nl solutions containing targeting or mismatch control morpholinos at a concentration of 7 μM each in nuclease free water, supplemented with 0.1% phenol red. Unlabelled and fluorescein-labelled morpholinos were ordered from Gene Tools LLC. Cy5 labelling of 3’alkyne tagged morpholino (obtained from Gene Tools LLC) was performed as detailed in Supplementary methods. At the low concentration used, MOs did not cause developmental defects in the majority of embryos, and did not affect mature miRNA processing from the primary transcripts (**Suppl. Fig. 1a,b**). Upon co-injection with mRuby3:H2B, mApple:PCNA fusion proteins or antibody the final concentration of proteins in the injection solution was 400 ng/μl for the fusion proteins and 250 ng/μl for the antibody. We verified that neither recombinant H2B protein nor injection of MO affected the cell cycle (**Suppl. Fig 3a,b**). After injection the embryos were kept in E3 media at 28°C until imaging or fixation.

### Transcription block

Transcription block was performed by treatment with 1 μM triptolide from the single-cell stage (Sigma T3652) in E3 media or by micro-injection of 200pg α-amanitin (Sigma A2263) at single cell stage.

### Live embryo imaging

Imaging of live embryos injected with fluorescent labelled morpholino oligonucleotides, fluorescent protein fusions or antibodies was performed on Zeiss Lightsheet Z1 or Zeiss 880 confocal microscope with Fast Airyscan Module.

For imaging on Zeiss Lightsheet Z1 embryos were mounted in 1% low melt agarose column using size 3 glass capillaries and incubated in E3 media at 28°C during imaging. Z-stacks of approximately 200-300 slices in 0.5-1 μm steps were acquired every 30-60 seconds for 1.5-2.5 hours, with 20x objective.

For imaging on the Zeiss 880 confocal microscope in Fast Airyscan mode embryos were mounted in 0.5% low melting agarose in glass bottom imaging dish and incubation chamber temperature was maintained at 28°C during imaging. Z-stacks of approximately 100-200 slices were acquired (step size of 0.25 μm) every 30-60 seconds for 1.5-2.5 hours using a 25x or 40x objectives. Post-processing of the acquired imageswas performed using Zeiss ZEN Blue Software.

### Analysis of spatio-temporal dynamics of MO signal

For global temporal analyses, lightsheet microscopy (morpholino/transgenic mRuby3:H2B) datasets were transformed into maximum intensity projections (MIPs), using Zeiss Zen Black software. Each MIP was imported into Icy Bioimage (Quantitative Image Analysis Unit, Pasteur Institute, France) where the spot detection plugin was used to detect transcription foci and nuclei, as these features are represented as regions of locally elevated intensity, in their respective channels. The plugin was guided to detect features as bright features against a darker background. Transcription foci were typically detected using a pixel size parameter of 3px, with a sensitivity of between 80 and 150% depending on noise (with higher sensitivity for higher noise). Feature size limits for detecting transcription foci were between 3 and 75 pixels in area. Nuclei were detected using a pixel size of between 13 and 25px, with similar sensitivity settings. Feature size limits for detecting nuclei were between 200 and 3000 pixels in area. Spot detection would be run on each channel of the image separately to detect the features being represented in that channel. Spot detection produced 2D regions of interest (ROIs) around defined features. Transcription foci were given a label which associated with their corresponding nucleus, and this was applied on a frame by frame basis to allow counting of nuclei expressing transcription foci. ROI metadata was exported to Microsoft Excel were transcription foci could be quantified.

### 3D analysis

3D datasets (Airyscan morpholino/injected mRuby3:H2B protein) collected from the Zeiss 880 confocal microscope with Fast Airyscan Module were imported into Icy Bioimage, where the spot detection plugin was used to identify nuclei and transcription foci, using the same sensitivity settings as described previously. Size features were scaled up to account for volumes: feature size settings for transcription foci were between 5-150 pixels, and size settings for nuclei were between 750 and 20,000 pixels. This would produce a 3D ROI around the detected feature. In the case of nuclei this ROI could be viewed in the 3D VTK viewer on Icy Bioimage, where a video of the rendered ROI could be recorded using the built-in recording function (Supplementary Figure 6). ROI data were exported to Microsoft Excel where transcription foci volumes were extracted per frame and timeseries could be plotted (Figure 5c).

In PCNA imaging data obtained on Zeiss 880 confocal with Airyscan, PCNA deficient regions were quantified using the spot detection plugin, with the setting to search for dark features against a bright background. Pixel sensitivity and 3d size settings were the same as for transcription foci detection, described previously. Relevant ROI’s were confirmed by visually inspecting that they correlated with signal deficient regions, and labelling the same feature with the same name per frame. The volume for each feature was extracted and plotted over time using Microsoft Excel.

### Cell cycle segmentation reference

Cell cycle phase segmentation was based on a visual library of cell cycle phase features generated from previously published confocal imaging of chromatin^65-67^. This reference library was used as a base for segmenting nuclei from MIP Zeiss Lightsheet data into cell cycle phases. Once phases were assigned to an initial batch of nuclei, the imaging parameters were extracted. These parameters were used to generate a quantitative measure of chromatin consistency throughout the cell cycle and assign a measure of cell cycle phase to each nucleus. Imaging parameters included 2D area covered by a nucleus ROI and standard deviation in pixel intensities across the nucleus (supplementary figure 3c).

### Whole mount antibody staining

Embryos were fixed at the desired stage in 4% PFA in PBS then permeabilised with PBS-0.3% TritonX-100. Blocking was performed with room temperature blocking solution (BlockAid™, B10710, Thermo Fisher) then transferred to blocking solution containing primary antibody. Excess primary antibody was removed with PBS-0.1% Tween-20 washes. Samples were then incubated with blocking solution containing appropriate secondary antibody conjugated to Alexa Fluor fluorescent dye at 1:500 dilution. Immunostained embryos were imaged in antifade mounting media with DAPI (Vectashield, H-1200, Vector Laboratories) on Zeiss 880 confocal microscope with Fast Airyscan Module.

Primary antibodies used:

- Anti-RNA polymerase II (phospho S2), ab5095, Abcam at 1:400 dilution
- Pol II S2p monoclonal antibody, C15200005, Diagenode at 1:400 dilution
- Anti-Histone H3 (di methyl K79) antibody, ab3594, Abcam at 1:1000 dilution

### Nascent RNA staining

Zebrafish embryos were injected with 1 nl of 50 mM ethynyl uridine (EU, Thermo Fisher, C10329) solution at single cell stage, incubated in E3 media at 28°C until fixation at the desired stage. Embryos were permeabilised with PBS-0.3% TritonX-100 before undergoing detection of nascent RNA using the Click-iT™ RNA Alexa Fluor™ 488 Imaging Kit (Thermo Fisher, C10329), following the manufacturer’s protocol. After the reaction embryos were imaged on Zeiss 880 confocal microscope with Fast Airyscan, or used for subsequent protein or DNA detection.

### DNA FISH

Probes for DNA-FISH were prepared using the FISH Tag™ DNA Multicolor Kit (Thermo Fisher) following kit instructions. The BAC DKEY-69C19 was used as a template for DNA-FISH probe production, corresponding to 214kb of the *mir430* locus on chromosome 4.

Embryos were fixed at the developmental stage of interest using 4% PFA. Animal caps were isolated and a series of gradient washes was used to equilibrate the samples with hybridization buffer [50% formamide, 4x SSC, 100mM NaPO4 pH 7.0, 0.1% Tween-20]. Embryonic DNA was denatured by incubation of samples at 70°C for 15 minutes before application of probe and incubation overnight at 37°C. Unbound probe was washed from the samples using a hybridization buffer: PBS-T gradient. Samples were counterstained with DAPI prior to imaging on a Zeiss LSM 880 with Fast Airyscan, with a 100 × 1.48 numerical aperture objective lens. 50-100 optical sections (130nm thickness) were acquired of each nucleus. Acquired images were processed using Zen software (Zeiss).

### Transgenic lines

The transgenic line Tg(Xla.Eef1a1:h2b-mRFP1) (https://zfin.org/ZDB-TGC0NSTRCT-130711-1 ^26^) used in this study was kindly provided by Sarah Baxendale (University of Sheffield, UK).

### Protein synthesis

mRuby3:H2B and mApple:PCNA were PCR amplified form plasmids *pKanCMV-mRuby3-10aa-H2B* (Addgene, 74258) and *mApple-PCNA-19-NLS-4* (Addgene, 54937) respectively and sub-cloned into NdeI/BamHI linearized *pET-28a+* plasmid (in frame with the N-terminal 6xHis-Tag), using In-Fusion® HD Cloning Plus Kit (Clontech, 638910) following the manufacture instructions.

The plasmids were transformed into *E. coli* BL21 (DE3) and the proteins were expressed in LB media by IPTG induction at 20°C. The cell lysates containing mApple:PCNA were prepared by re-suspending in 50 mM Tris HCl; pH8, 250 mM NaCl, 10 mM imidazole, 2 mM MgCl2 and 10% glycerol whereas mRuby3:H2B was resuspended in 50 mM Tris HCl; pH 8, 500 mM NaCl, 10 mM imidazole, 2m M MgCl2, 5% glycerol and 0.3% Brij. All buffers were additionally supplemented with BitNuclease, PMSF and protease inhibitors. Cell lysis was improved by subjecting the lysates to sonication followed by centrifugation to clarify the cell extracts. The proteins were then purified by Nickel affinity chromatography (Generon). The proteins were dialysed against 500 mM KCl, 20 mM PIPES, 100 uM EGTA, pH6.8 using 12 kDa columns (Pur-A-Lyzer; Sigma).

### Manipulation of embryonic cell ploidy by heat-shock treatment

Tetraploid embryos were generated by heat-shock treatment, following the HS2 protocol adapted from^44^. Embryos were collected 2 minutes post fertilization and maintained in E3 media for a further 20 minutes at 28°C, during which microinjection of morpholino oligonucleotides was performed. Embryos were heat-shocked between 20-23 minutes post fertilization by incubation in E3 media at 42°C for 2 minutes, then returned to E3 media at 28°C and allowed to develop normally. Embryos were monitored at the normal point of the second cell division (1 hour post fertilisation (hpf)) and heat-shock treated embryos that failed to undergo cytokinesis were selected for downstream analysis, along with matched non-heat-shock treated controls. For the assessment of ploidy manipulation, embryos were collected at the 12-somite stage, dissociated with PBS-based enzyme-free cell dissociation buffer (Gibco), washed with PBS and subjected to propidium iodide DNA content analysis following manufacturer’s conditions (Invitrogen).

### Cell cycle length and gene expression experiments

Wild type embryos were injected with mRuby3:H2B protein at the single cell stage and incubated in E3 media supplemented with 1μM triptolide (Sigma, T3652) or vehicle control following injection. In cell cycle elongation experiments, wild type embryos were injected at the single cell stage with a cocktail of 200pg of *Xenopus chk1* mRNA, 200pg miR430 MO and mRuby3:H2B protein whereas controls were injected with 200pg *mRuby3* mRNA + 200pg miR430 MO and mRuby3:H2B protein. Embryos were imaged using Zeiss Lightsheet Z.1 microscope with the ZEN software. Images were analysed and timing of cell cycle transition from anaphase to anaphase were identified by mRuby3:H2B pattern variation using Icy Bioimage. Statistical analysis was carried out by Prism GraphPad 5 (USA). *Chk1* mRNA and *mRuby3* mRNA were produced from linearised pCS2+ vector by mMESSAGE mMACHINE SP6 Transcription Kit (ThermoFisher Scientific).

### Quantification of *miR430a* and *miR430b* levels by digital droplet PCR (ddPCR)

Total RNA was extracted from a 100 miR430 targeting MO1-3 and non- injected embryos at sphere stage, using TRIzol™ Reagent (ThermoFisher, 15596026) flowing manufacturer's instructions. Three RNA samples for each group (MOs injected and noninjected) from three independent experiments were prepared. cDNA was synthesized from equal amounts of total RNA (100ng) using the High-Capacity cDNA Reverse Transcription Kit (Applied Biosystems) following the manufacturer’s protocol. cDNA samples were diluted 1:30 with nuclease-free water. qPCR reactions were prepared using the ddSupermix for Probes (Bio-rad) and Taqman miRNA probes miR430a (004365_mat) and miR430b (465767_mat). Droplet generation oil was added to the qPCR mixture and droplets were generated with the QX100 droplet generator (Bio-rad). PCR protocol as per manufacturer (Taqman miRNA assay) and the samples were run in a QX100 droplet reader. Quantasoft software (Bio-rad) was used for data acquisition and analysis.

